# Deep Neural Network for Protein Contact Prediction by Weighting Sequences in a Multiple Sequence Alignment

**DOI:** 10.1101/331926

**Authors:** Hiroyuki Fukuda, Kentaro Tomii

## Abstract

Protein contact prediction is a crucially important step for protein structure prediction. To predict a contact, approaches of two types are used: evolutionary coupling analysis (ECA) and supervised learning. ECA uses a large multiple sequence alignment (MSA) of homologue sequences and extract correlation information between residues. Supervised learning uses ECA analysis results as input features and can produce higher accuracy. As described herein, we present a new approach to contact prediction which can both extract correlation information and predict contacts in a supervised manner directly from MSA using a deep neural network (DNN). Using DNN, we can obtain higher accuracy than with earlier ECA methods. Simultaneously, we can weight each sequence in MSA to eliminate noise sequences automatically in a supervised way. It is expected that the combination of our method and other meta-learning methods can provide much higher accuracy of contact prediction.

## 1. Introduction

Protein contact prediction is a crucially important step for protein structure prediction. Many contact prediction methods have been developed [1–19]. In earlier stages of contact prediction history, most successful prediction methods were based on evolutionary coupling analysis (ECA) of large multiple sequence alignment (MSA) of homologue sequences. In evolutionary processes, pairs of residues that are near each other in the tertiary structure reportedly tend to co-evolve to maintain their structure. For instance, when one becomes larger, the other becomes smaller. Alternatively, when one becomes a positively charged amino acid, the other becomes a negatively charged amino acid.

Usually evolutionary information has many noise signals because indirect correlation can be seen observed between residues (A and B) when residues (A and C) and residues (B and C) are correlated. We must extract true correlation from such noise. Many challenges have arisen in this field. The methods used to address them can be categorized into two groups: Graphical Lasso and pseudo-likelihood maximization. Graphical Lasso, a graph structure estimation method, was developed by Friedman et al. in 2008 [20]. It can estimate the graph structure from a covariance matrix using likelihood estimation of a precision matrix with L1 regularization. PSICOV [4] and EVFold [5] are famous programs that apply Graphical Lasso to contact prediction problems. However, pseudo-likelihood method is an approximation method for probabilistic models such as a Potts model to estimate the interaction strength between residues. It is usually difficult to calculate the marginal probability exactly. Therefore, such an approximation method is often used. Major programs using this method are plmDCA [11], GREMLIN [7], and CCMpred [13].

After these extensive studies of ECA, meta-supervised learning methods emerged for protein contact prediction using the results of these ECA methods as input features. MetaPSICOV [14], a well-known supervised method, uses the outputs of PSICOV, CCMpred, and FreeContact [12] as input features and uses other features such as secondary structure probability, solvent accessibility, and Shannon entropy. In all, 672 features were used. MetaPSICOV succeeded in improving prediction accuracy much more than a single ECA method. Recently, Wang et al. [15] developed an ultra Deep Neural Network method and achieved state-of-the-art of prediction accuracy.

However, in spite of recent success obtained using these supervised methods, these methods require ECA results. Moreover, they do not directly predict contacts from MSA. As presented herein, we describe a novel approach to contact prediction that can both extract correlation information and predict contacts in a supervised manner directly from MSA using a deep neural network (DNN). Using DNN, we can predict contacts more accurately than previous ECA methods can. Furthermore, computation takes much less time because no iterative calculation is necessary once the training process has been completed.

Furthermore, our DNN-based method can provide a means of optimizing the input MSAs in a supervised manner: The weight of each sequence in MSA is parameterized. It can be optimized through DNN so that we can eliminate noise sequences in MSA automatically. In this model, we expect that more important sequences have greater weight and that less important sequences have less weight after the optimization. Today, a growing number of protein sequences are obtainable so that not all sequences in MSA necessarily have the same contacts. These sequences result in noises for contact prediction. In addition, Fox et al. [21] have shown that the accuracy of contact prediction depends on the MSA accuracy. Driven by such a motivation, we also try to weight the sequences of MSA correctly.

## 2. Materials and Methods

### 2.1 Dataset

An original dataset was prepared for this experiment using the following steps. 1) A set of non-redundant amino acid sequences was obtained from the PISCES cull pdb server (sequence identity cutoff, 20%; resolution cutoff, 2.5 Å; R-factor cutoff, 1.0; generated day, July 20, 2017; number of chains, 12094) [22]. 2) PDB files were retrieved and true contact pairs were calculated from protein backbone coordinates. In this experiment, we infer contact if the distance of C_β_ atoms of the residue pair is less than 8 Å. For glycine residues, C_α_ atoms were used instead of C_β_ atoms. PDB coordinates include many missing values (in our dataset, more than 4000 proteins have at least one missing value about C_β_ atoms). Therefore, we marked a residue pair which had a missing C_β_ coordinate as NaN and excluded it when we calculated the loss. 3) Removal of redundancy with the test set was performed. We excluded training proteins sharing >25% sequence identity or having BLAST E-value <0.1 with any test protein by blastp. 4) Proteins with length of greater than 700 residues and with fewer 25 residues were also eliminated. Finally, our original dataset consists of 10038 proteins. We split them into 9038 proteins (training set) and 1000 proteins (validation set). As the test sets, for benchmarking our method, we used the dataset that was used in the PSICOV paper [4] and the CASP11 dataset [24]. 5) The MSAs for proteins in datasets were obtained using HHBlits [23] with three iterations. The E-value was set to 0.001 on the UniProt20_2016 library.

### 2.2 Model

Our challenge is to predict contact directly from correlation information using DNN. Similarly to Graphical Lasso methods, we take covariance matrices calculated from MSA as input. Then we feed them into the neural network to obtain the probability of the contact. For the calculation of covariance matrices, we used the same formula as that of the PSICOV paper as shown below.

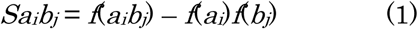

Therein, *a* and *b* are amino acid types at positions *i* and *j*, respectively, *f*(*a*_*i*_) (and *f*(*b*_*j*_)) is a frequency of amino acid *a* (and *b*) at position *i* (and *j*), and *f*(*a*_*i*_*b*_*j*_) is a frequency of amino acid pairs *a* and *b* at position *i* and *j*. If no correlation is observed between *i* and *j* with respect to amino acid pairs *a* and *b*, then *Sa*_*i*_*b*_*j*_ is equal to zero. By this formula, with pairs of 21 amino acid type (including gap), one can obtain 441 of L×L covariance matrices where *L* is the sequence length of the target protein.

Here, our network architecture is presented in Fig. 1(A). We are inspired by a study reported by Belilovsky et al. [25] showing that DNN is applicable to graph estimation problems. For that reason, we chose to apply DNN to contact prediction. Our input covariance matrices are regarded as L×L pixel images with 441 channels (typical color images have three channels). Therefore, we can apply a convolutional neural network (CNN), which is widely applied on image classification tasks. For this experiment, we introduce dilated CNN, a kind of CNN used at semantic segmentation tasks, because we must predict the contact on each pixel and because it is similar to the situation of segmentation tasks. In addition, because proteins usually have long-range interactions and because dilated CNN can cover a wider range of interaction using the same number of parameters as normal CNN, it is beneficial for contact prediction. Although many architectures of network can be presented today, we first use the one presented in the original paper using dilated CNN on semantic segmentation tasks [26]. The feature maps of our network are presented in Fig. 1(C).

**Fig. 1.**
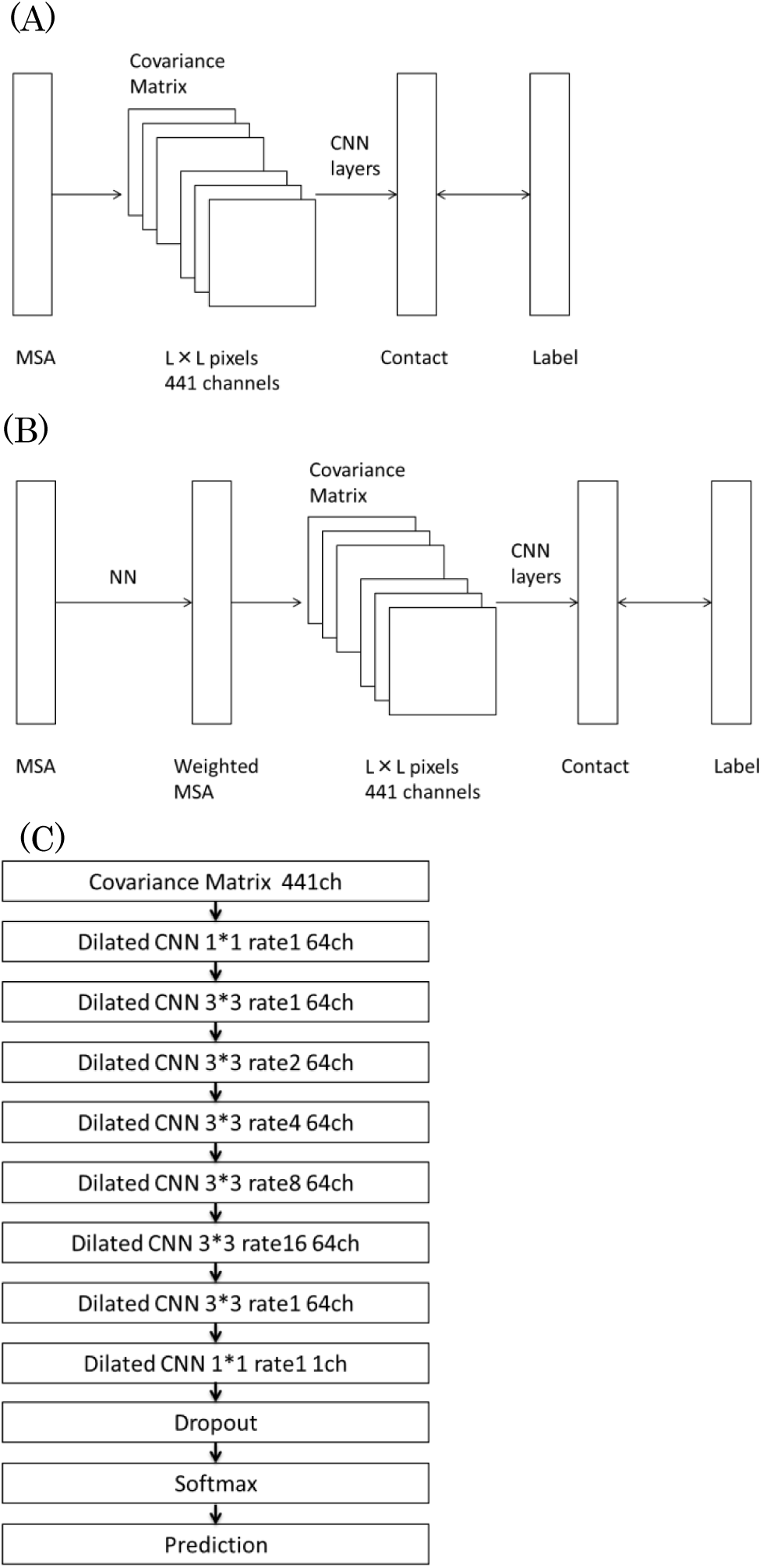
(A) Dilated CNN used in our experiments. (B) MLP is connected to the dilated CNN through the weighted MSA. (C) Exact architecture of our dilated CNN.

Additionally, to eliminate noise of MSA, we weight each sequence of MSA. This weighting is also performed using neural networks. We built a multilayer perceptron network that outputs the weight of each sequence in MSA. This network has seven features as input: number of sequences in an alignment, sequence identity with a query sequence, sequence identity with a consensus sequence, gap ratio, and averages of the last three features. Our aim is that less important sequences that have many gaps are less weighted. Multilayer perceptrons (MLPs), which have three hidden layers and for which each hidden layer has seven nodes, are used. The output of this network is used to weight MSA. From weighted MSA, 441 covariance matrices are calculated and are fed to dilated CNN as input. We can optimize the network completely. In other words, we can completely learn both parameters of how to weight the MSA and how to predict contact. Our total network is presented in Fig. 1(B).

We trained these networks using the ADAM optimizer with the learning rate of 0.0005. Batch Normalization is used to obtain higher accuracy and faster convergence. Dropout is used to avoid overfitting. All proteins have different lengths. All input matrices have different sizes. Therefore, we calculate the graph by one batch size. All hyperparameters and network architectures are selected by the score of validation set. All experiments were conducted on the ordinary desktop computer with a GPU (GeForce TITAN X; Nvidia Corp.) using the TensorFlow library. Training required several days to calculate 20–30 epochs.

### 2.3 Evaluation

Evaluation of our method was done by comparison or results with those obtained using existing methods. We define the contact distances as “short” 6<=|*i*-*j*|<=11, “medium” 12<=|*i*-*j*|<=23, and “long” 24<=|*i*-*j*|, and evaluate top *L*/*k* (*k*=10,5,2,1) prediction. The prediction accuracy was calculated using the following equation.

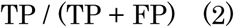

In that equation, TP denotes the number of true contacts among the predicted ones; TP + FP is the number of all predicted contacts. We selected PSICOV, CCMpred, and MetaPSICOV as existing methods to be compared and calculated with local prediction by ourselves directed by instructions for using each method. Using the CASP11 dataset, we calculated the accuracy for each separated domain, not a whole protein.

## 3. Resultsand Discussion

### 3.1 PSICOV dataset

Table 1 presents the predictive accuracy of three existing methods and our method. Our method outperformed the existing ECA methods at every distance and prediction count. Although we use only correlation information based on an MSA and do not use any other feature such a secondary structure, our method outperformed the meta-learning method: metaPSICOV.

**Table 1.** Contact prediction accuracy with the PSICOV dataset. Bold characters signify the highest accuracy within the column.

| Method | Short |  |  |  | Medium |  |  |  | Long |  |  |  |
| --- | --- | --- | --- | --- | --- | --- | --- | --- | --- | --- | --- | --- |
|  | L/10 | L/5 | L/2 | L | L/10 | L/5 | L/2 | L | L/10 | L/5 | L/2 | L |
| PSICOV | 0.55 | 0.42 | 0.25 | 0.16 | 0.63 | 0.51 | 0.31 | 0.2 | 0.77 | 0.69 | 0.53 | 0.37 |
| CCMpred | 0.65 | 0.5 | 0.29 | 0.19 | 0.73 | 0.6 | 0.36 | 0.23 | 0.8 | 0.75 | 0.62 | 0.45 |
| DNN | 0.84 | 0.72 | 0.47 | 0.28 | 0.84 | 0.73 | 0.52 | 0.34 | **0.92** | 0.87 | 0.75 | 0.58 |
| DNN+weightedMSA | **0.86** | **0.74** | **0.48** | **0.29** | **0.85** | **0.76** | **0.53** | **0.35** | **0.92** | **0.88** | **0.76** | **0.6** |
| MetaPSICOV | 0.82 | 0.7 | 0.46 | 0.27 | 0.82 | 0.72 | 0.51 | 0.33 | 0.9 | 0.86 | 0.72 | 0.57 |

When extracting correlation information for one residue pair, 21 × 21 correlation scores from 21 × 21 amino acid pairs are obtained. However, these scores are merely averaged in CCMpred and PSICOV. By contrast, our method uses 441 covariance matrices as input features and feeds them to the CNN architecture. This does not lead the loss of information, which is an important benefit of our method compared to PSICOV and CCMpred. Moreover, the CNN architecture can extract useful features from covariance matrices automatically through convolutional operation.

### 3.2 CASP11 dataset

We also evaluate our method using the CASP11 dataset, which is more difficult to predict because almost all CASP11 proteins have an insufficient number of homologue sequences. Table 2 shows the results. As expected, the difference between DNN and DNN+weightedMSA is smaller than that obtained in the case of using the PSICOV dataset. However, even for a harder task, our method can perform well. In the case of the CASP11 dataset, MetaPSICOV has greater accuracy of short-range prediction, although our method performs well at a long range by a large margin which is more important to predict the tertiary structure.

**Table 2.** Contact Prediction Accuracy on CASP11 dataset. Bold characters show the highest accuracy within the column.

| Method | Short |  |  |  |  | Medium |  |  |  |  | Long |  |  |  |
| --- | --- | --- | --- | --- | --- | --- | --- | --- | --- | --- | --- | --- | --- | --- |
|  | L/10 | L/5 | L/2 | L |  | L/10 | L/5 | L/2 | L |  | L/10 | L/5 | L/2 | L |
| PSICOV | 0.32 | 0.24 | 0.16 | 0.12 |  | 0.35 | 0.27 | 0.19 | 0.13 |  | 0.4 | 0.35 | 0.26 | 0.2 |
| CCMpred | 0.36 | 0.28 | 0.18 | 0.13 |  | 0.41 | 0.32 | 0.22 | 0.15 |  | 0.45 | 0.41 | 0.32 | 0.24 |
| DNN | 0.67 | 0.56 | 0.38 | 0.24 |  | **0.69** | **0.59** | **0.43** | **0.29** |  | **0.68** | 0.63 | 0.53 | 0.41 |
| DNN+weightedMSA | 0.66 | 0.56 | 0.38 | **0.25** |  | 0.67 | **0.59** | **0.43** | **0.29** |  | **0.68** | **0.65** | **0.54** | **0.42** |
| MetaPSICOV | **0.71** | **0.59** | **0.4** | **0.25** |  | **0.69** | **0.59** | 0.42 | **0.29** |  | 0.58 | 0.55 | 0.46 | 0.36 |

### 3.3 Effects of Weighting the MSA

Here, we demonstrate that weighting of the MSA can boost the prediction accuracy. Our network can correctly learn how to weight the MSA sequence. Figure 2(A) shows the distribution of weight values of one protein. We can find that these were well distributed. Some values were nearly zero, which shows that some noise sequences existed in the original MSA.

**Fig. 2.**
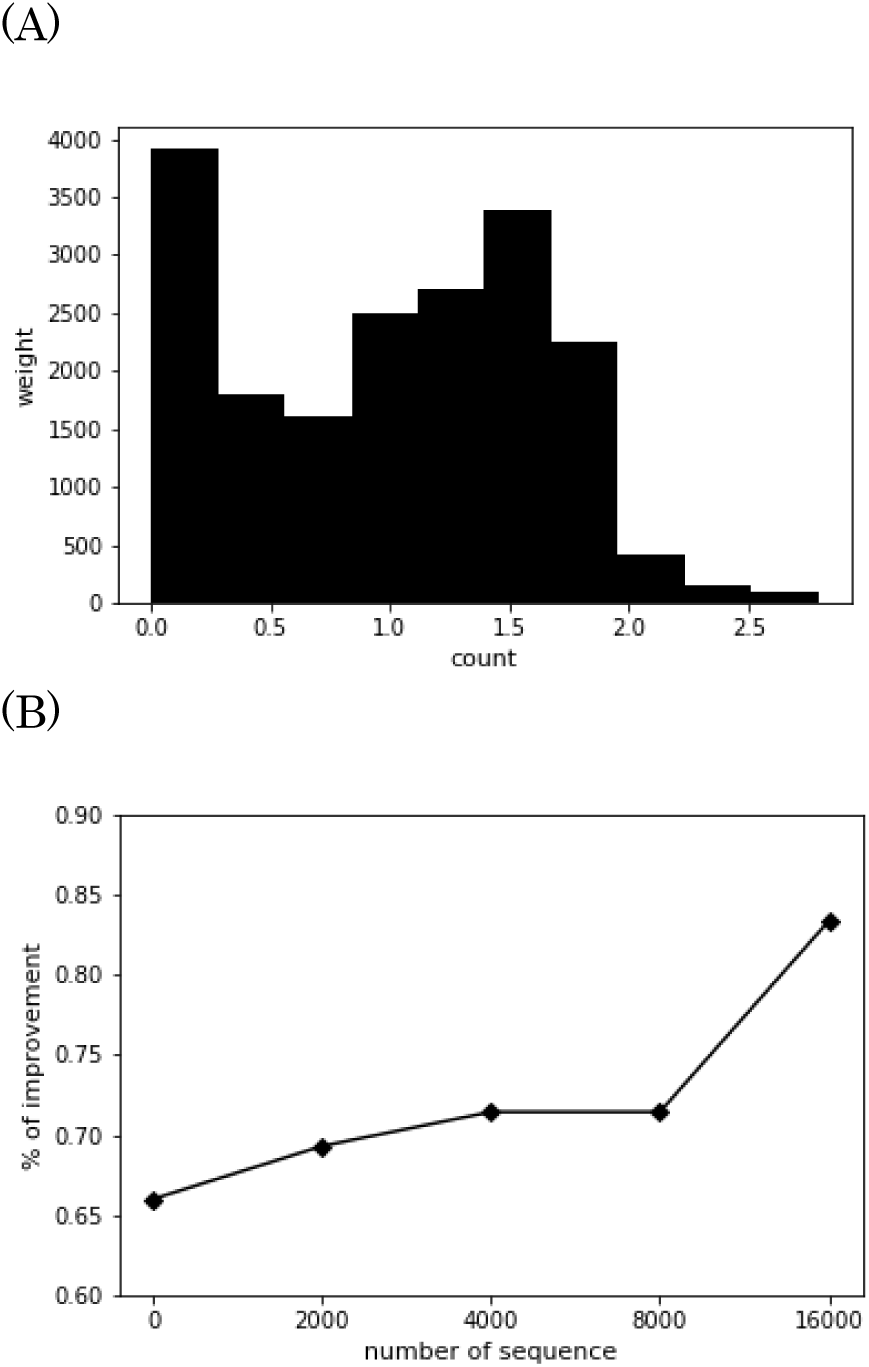
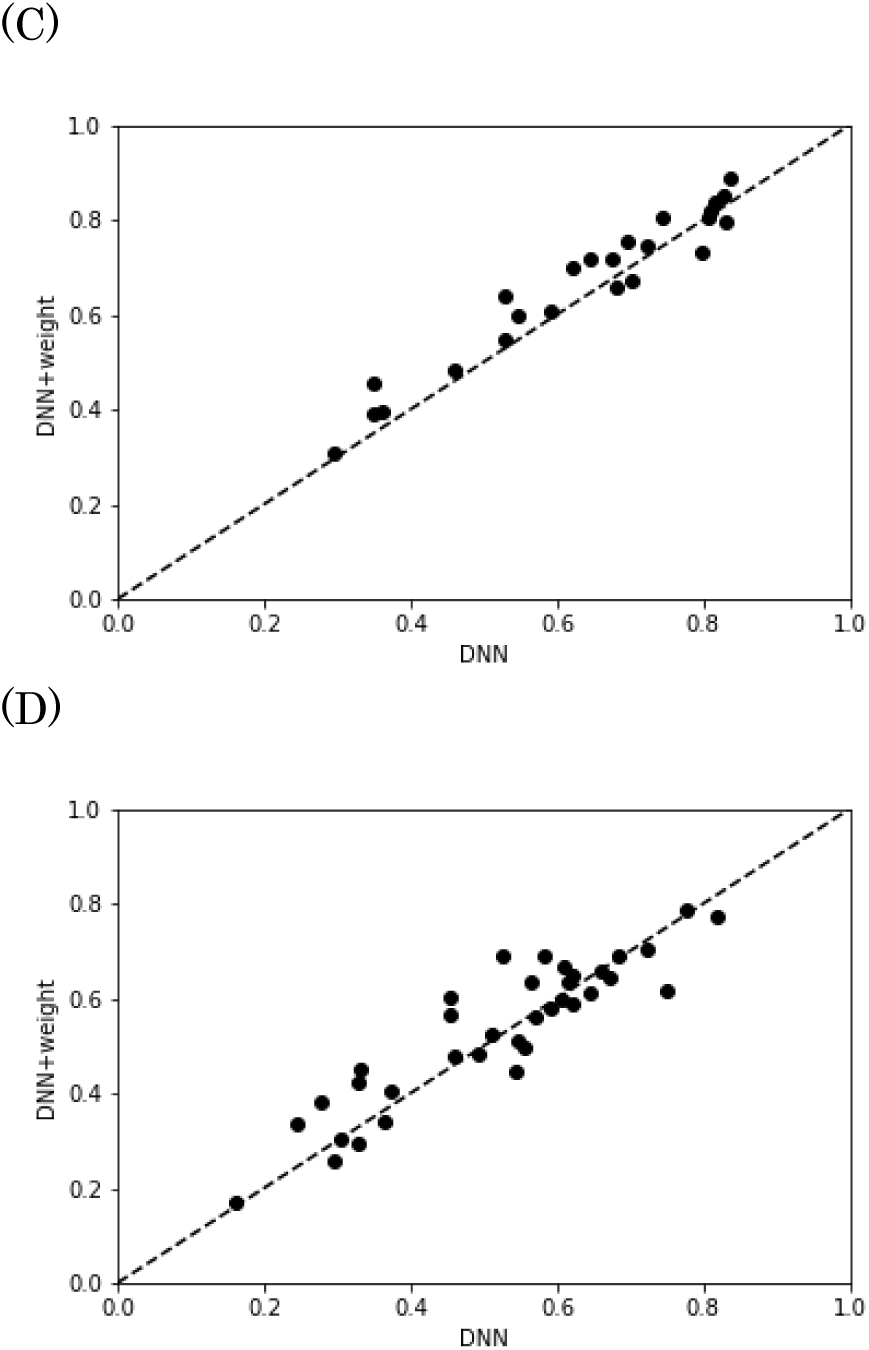
(A) One example of weight distribution in the sequences of one MSA. (B) The dependence of improvement of accuracy on the number of sequences. (C) DNN method top L accuracy shown against the DNN+weight method when we have over 16,000 homologue sequences. (D) shows that of fewer than 2000 homologue sequences.

To investigate the result further, we calculate the dependence of prediction accuracy on the number of sequences in MSA. We use the top L prediction at long range to compare the accuracy on the PSICOV dataset because this area has the greatest number of predictions and because the standard deviation is smallest. Figure 2(B) shows that when we have numerous sequences, we can improve the prediction accuracy, but when we have only a few sequences, we cannot improve it. The percentage of improvement is the number of improved proteins divided by the total number of proteins. This result demonstrates that the network can remove noise sequences when MSA has a sufficient number of homologue sequences.

### 3.4 Calculation Time

In terms of calculation time, our method also shows good performance. Here, we compare the calculation time of our method with CCMpred, which is the fastest method among existing ECA methods. Table 3 shows that our method takes much less time than the CCMpred with and without GPU. Although Graphical Lasso and pseudo-likelihood method have iterative calculations, neural network methods can calculate the result directly. We can obtain results in a short time once we have completed the network training. Our method is practically useful when huge numbers of contact predictions are needed.

**Table 3.**

|  | CCMpred (CPU) | CCMpred (GPU) | Our DNN (CPU) | Our DNN (GPU) |
| --- | --- | --- | --- | --- |
| Time (min) | 585 | 47 | 10 | 2 |

## 4. Conclusions

As described in this paper, we present the DNN approach to contact prediction and weighting MSA. Our method performs better than any other ECA method. Moreover, it can obtain comparable results to those of MetaPSICOV, although we use only correlation information based on MSA as an input. Future studies can use our results as input features of other meta-learning methods or can add any other feature such as secondary structure information to our method. Although Graphical Lasso and pseudo-likelihood methods are independent of the number of data available, our method is a supervised method that depends on the dataset size. Today, because of numerous protein structural data that have become available through extensive efforts, supervised learning will gain accuracy.

Recently, DNN-based methods have succeeded in various fields. We think DNN can provide many benefits for contact prediction. 1) It can predict the whole protein contacts at once, whereas other supervised methods such as RandomForest or SVM predict only one pair of residues. 2) Variable lengths of proteins are treated uniformly. 3) Using back propagation, end-to-end learning can be performed. By virtue of this advantage, DNN provides a chance to optimize contact prediction and refinement of MSA simultaneously in a supervised way. As the next step, using DNN, we can try network architectures of various kinds extensively in the hopes of obtaining further improvement of prediction accuracy.

## Additional Note

During our preparation of this paper, another paper which reports contact prediction method applying DNN directly to MSA is published. [27] However, our research is done by ourselves independently.

## ACKNOWLEDGEMENTS

Computations were performed partially on the NIG supercomputer at ROIS National Institute of Genetics. We deeply thank the National Institute of Genetics for supporting our research.

This research was partially supported by Platform Project for Supporting Drug Discovery and Life Science Research (Basis for Supporting Innovative Drug Discovery and Life Science Research (BINDS)) from AMED under Grant Number JP17am0101110.

